# The effect of stress on the balance between goal-directed and habit networks in obsessive-compulsive disorder

**DOI:** 10.1101/598268

**Authors:** Anouk van der Straten, Wieke van Leeuwen, Damiaan Denys, Hein van Marle, Guido van Wingen

**Author notes:** **Contact information:** Anouk van der Straten, MD, Department of Psychiatry, Amsterdam UMC, University of Amsterdam, Meibergdreef 5, 1105 AZ Amsterdam, The Netherlands.

## Abstract

The classical cognitive-behavioral theory of obsessive-compulsive disorder (OCD) holds that compulsions are performed to reduce distress that is evoked by obsessions, whereas a recent neuroscience-inspired theory suggests that compulsivity results from a disbalance between goal-directed and habit-related neural networks. To bridge these theories, we investigated whether the balance between goal-directed and habit networks in patients with OCD was affected in the late aftermath of stress. Twenty-three OCD patients and twenty-three healthy controls participated in a controlled stress induction paradigm using the socially evaluated cold-pressor test in a crossover design. Stress responses were evaluated through cortisol levels, blood pressure, and anxiety ratings. Functional connectivity of the caudate nucleus and posterior putamen was assessed using seed region analysis of resting-state functional magnetic resonance imaging data, which are hubs of the goal-directed and habit network, respectively. Stress induction increased blood pressure and psychological stress measures across groups and resulted in blunted cortisol responses in patients. Furthermore, patients showed a blunted reduction in connectivity between the caudate nucleus and precuneus in the aftermath of stress, which was positively correlated with compulsivity but not obsession severity. The posterior putamen showed no significant group differences in stress-induced connectivity. These results suggest that compulsivity in OCD in the aftermath of stress is associated with altered connectivity between the goal-directed and default mode networks.

## Introduction

Obsessive-compulsive disorder (OCD) is a debilitating psychiatric disorder with an estimated lifetime prevalence of 1-3% [1]. Patients with OCD suffer from obsessions, defined by recurrent intrusive anxious thoughts or images, and/or compulsions, defined by repetitive behaviors or mental acts. Although the patients are usually aware of the senselessness of the compulsions they are performing, they are unable to inhibit their behaviors. Currently, two apparently opposing theories have been postulated to explain compulsivity in OCD. The classic cognitive-behavioral theory proposes that performing compulsions is primarily a strategy to reduce distress that is evoked by obsessions [2]. This theory provides an explanation of OCD symptoms that forms the basis of current cognitive-behavioral therapy. However, the variety in phenomenology shows that the relationship between obsessionality and compulsivity is more complex [3]. Recent neuroscientific studies suggest that patients with OCD show a general impairment in goal-directed behavior, resulting in an overreliance on habits [4,5]. Goal-directed behavior consists of actions that are performed to achieve a desired goal. When actions are performed on a regular basis, habits are formed to facilitate actions that do not require planning or organization. This leads to greater efficiency, but often at the cost of behavioral flexibility. Along these lines, the overreliance on (maladaptive) habitual behavior is thought to contribute to the development of compulsivity in OCD [6].

Both rodent and human studies have pointed towards the involvement of the posterior putamen and the caudate nucleus in the habit and goal-directed network, respectively [6-10]. Both structures are part of the corticostriatal circuitry, the dysfunction of which is seen as the neuroanatomical basis of OCD [11,12]. While the posterior putamen mediates habitual actions by targeting motor areas including the supplementary motor area (SMA), goal directed behavior is mediated by the interaction between the ventral medial prefrontal cortex (vmPFC) and the caudate nucleus [9]. The recent focus on the habit hypothesis became apparent in the discussion around the new DSM-5, in which OCD moved out of the chapter ‘anxiety disorders’ to obtain its own chapter: i.e. ‘obsessive–compulsive and related disorders’ [13]. The habit theory however, does not account for the typical clinical observation that psychological stress exacerbates OCD symptoms [14]. Furthermore, recent studies in healthy humans have shown that stress reduces prefrontal cortex-related goal-directed control and thereby induces a bias towards reliance on habitual behavior [15,16]. Integrating these findings leads to the hypothesis that stress may reduce goal-directed control and induce a shift towards habits, thereby contributing to the compulsivity seen in OCD [6,17]. However, this novel model that connects the critical role of stress as posited by the classical cognitive-behavioral theory with the neuroscience-inspired habit theory of OCD remains to be tested.

In this study, we investigated the late effects of stress on resting-state functional connectivity of both the caudate nucleus (as major hub within the goal-directed network), and the posterior putamen (as major hub of the habit network) comparing OCD patients with matched healthy controls. The effects of acute stress vary over time due to the rapid release of neuromodulators and the delayed release of corticosteroids, In addition, corticosteroids also have time-dependent effects related to rapid non-genomic and slow non-genomic effects [18]. As a result, catecholaminergic and dopaminergic systems are activated directly after a stressor, while the slower genomic effects of corticosteroids can continue for several hours and affect cognitive and emotional processing [19-21]. We hypothesized that in OCD patients, connectivity is reduced within the goal-directed network (i.e. between the caudate nucleus and the ventromedial prefrontal cortex) and/or increased within the habit network (i.e. between the posterior putamen and supplementary motor area) in the late aftermath of stress.

## Methods and Materials

### Participants

Twenty-five OCD patients were recruited from the outpatient clinic for anxiety disorders at the psychiatry department of the Amsterdam UMC and through advertisement at a Dutch patient organization website. Inclusion criteria were (1) being aged 18-65 years; (2) having a diagnosis of OCD according to the DSM-IV; (3) having a Yale-Brown Obsessive-Compulsive Scale (Y-BOCS) score of 12 or higher; (4) having provided written informed consent and being willing and able to understand, participate and comply with the study requirements. Exclusion criteria are reported in the Supporting Information. In addition, twenty-five healthy controls without a current or past psychiatric diagnosis as assessed with the M.I.N.I. and matched according to age, sex, and educational level, were recruited through flyers and online advertisements. All participants were assessed on anxiety, depressive and obsessive-compulsive symptoms, using the Hamilton Anxiety Rating Scale (HAM-A) [22], the Hamilton Depression Rating Scale (HDRS) [23] and the YBOCS [24], respectively. In addition, the presence of psychiatric co-morbidities was assessed with the M.I.N.I. [25] and the Structured Interview for DSM-IV Personality (SIDP-IV) [26]. This study was approved by the Ethical Committee of the Academic Medical Center in Amsterdam (METC 2014_168) and all participants provided written informed consent. The study was registered in the ISRCTN registry (ISRCTN47698087).

### Design and measurements

The study used a cross-over design with counterbalanced order of stress induction vs. neutral condition across participants, with the two sessions separated by a 1 week interval. All the sessions took place in the afternoon to control for the normal daily fluctuations of cortisol. During the stress session, participants were exposed to the socially evaluated cold-pressor test (SECPT). This is a relatively short, extensively studied procedure which is known to induce a reliable physiological and subjective stress response through a minimally invasive method [27] (see Supplemental Information). The cold stress procedure causes vasoconstriction, which consequently leads to increased blood pressure and heart rate deceleration [27]. The physiological stress response was examined by measuring blood pressure and heart rate at five time points: at baseline, once during the water immersion procedure, after the math task, before the scanning session (45 minutes before the resting-state scan), and after the scanning session (40 minutes after the resting-state scan), see Figure 1. In addition, cortisol was measured through saliva sampling at four time points: at baseline, after the math task, right before the resting-state scan, and after the scanning session. Moreover, to assess the subjective stress response, participants were asked to rate their anxiety level by filling out the State-Trait Anxiety Inventory (STAI [28]) at four time points: prior to the water procedure, after the math task, right before the resting-state scan, and after the scanning session. Furthermore, participants had to rate how difficult, unpleasant, and stressful they experienced the stress procedure right after the math task using a subjective stress level questionnaire with a VAS scale [27].

**Figure 1.**
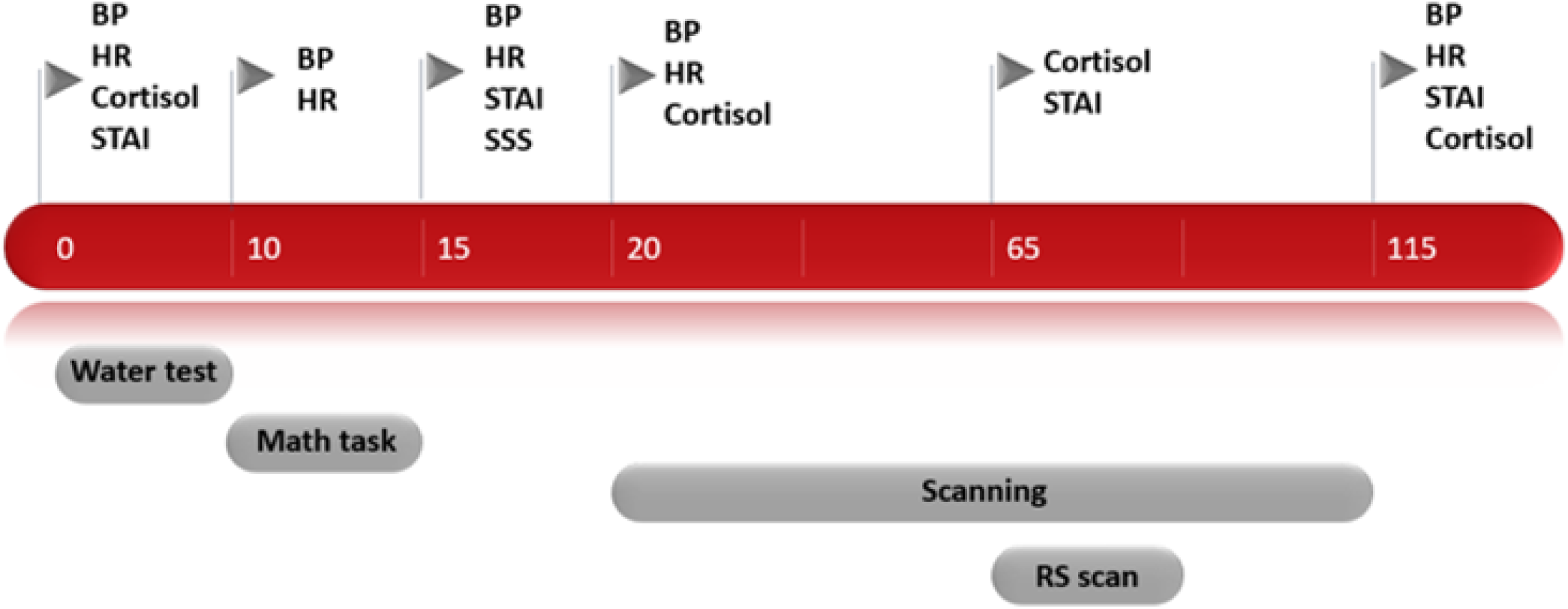
Timeline of experimental procedure and measurements. *Abbreviations: blood pressure (BP), heart rate (HR), State-Trait Anxiety Inventory (STAI), subjective stress scale (SSS) and resting-state (RS)S) scan*.

Because this study was part of a larger set of experiments, the resting-state scan was conducted approximately 65 minutes after the stress induction procedure. At the end of the stress session there was a short debriefing, in which the participants were told that the purpose of the test was to create stress. For an overview of the study design and measurements, see Figure 1.

### Data acquisition

MR imaging was performed on a 3.0 LT Philips MRI scanner (Philips, Best, The Netherlands), using a 32-channel SENSE head coil. The scanning protocol (see Supplemental Information) included a high-resolution T1-weighted MRI (voxel size=1.1mm isotropic). For fMRI, multi-echo echoplanar imaging (EPI) was used to acquire T2*-weighted MRI volumes with blood oxygen level-dependent (BOLD) contrast (repetition time=2375ms; voxel size=3.0 Lmm isotropic). Multi-echo EPI reduces signal dropout and distortion while enhancing BOLD contrast sensitivity [29]. Participants were instructed to stay awake and relax with their eyes open while thinking of anything that came to their mind.

### Data analysis

Demographical and clinical data were analysed with SPSS software (version 24.0, Chicago, IL, USA). Group differences were tested using Mann-Whitney U tests for continues not normally distributed variables (age, Y-BOCS score, HAM-A, HDRS, mean FD) and χ2 tests for gender, handedness and education. The physiological stress measures (blood pressure, heart rate and cortisol) and anxiety levels (STAI) were analysed using mixed model analysis of variance (ANOVA) with the factors group (patients, healthy controls), condition (stress, control) and time (five time points for blood pressure and heart rate, four for anxiety), corrected for the order of sessions. No time factor was used for cortisol, as we calculated the area under the curve with respect to the ground (AUCg) to assess overall exposure to cortisol during the session [30]. We additionally compared baseline cortisol values between the conditions to ensure that the results were not due to significant differences at baseline. Baseline cortisol and AUCg values were log-transformed before testing because of a non-normal distribution. Each question of the subjective stress level questionnaire was entered in a mixed model ANOVA with the factors factors group (patients, healthy controls), condition (stress, control) and question type (how difficult, how stressful, how unpleasant), corrected for the order of sessions. When the assumption of sphericity was violated the ANOVA results were corrected with a Greenhouse Geisser correction. The analyses were followed by post-hoc paired-t-testing in case of significance, with Bonferroni correction for multiple comparisons.

### MRI analysis

Standard preprocessing of functional MRI data was conducted in SPM12 (www.fil.ion.ucl.ac.uk/spm/) and the CONN toolbox v18 (http://www.nitrc.org/projects/conn) including realignment, slice-time correction, normalization to Montreal Neurological Institute (MNI) space and 6mm smoothing. Participants with realignment parameters exceeding 3mm translation on the x-, y-, or z-axis were excluded from the analysis, resulting in the exclusion of two patients. One healthy control was excluded due to an incidental neurological finding and one control was excluded because of insufficient brain coverage due to inaccurate setting of the field of view. This led to the inclusion of twenty-three patients and twenty-three healthy controls for further analysis.

We assessed functional connectivity of the left and right caudate and left and right posterior putamen using a seed-to-voxel analysis to investigate whether there were group differences in stress-induced connectivity. Data were corrected for nuisance variables using CompCor [31], band-pass filtered, and transformed using Fisher’s r-to-z transformation [32]. Because the caudate nucleus and posterior putamen are located closely to each other, the timeseries from the ipsilateral seed were regressed out to ensure that the results were specific for the selected seeds and did not reflect striatal connectivity in general. Connectivity maps were entered into a second-level group (patient, healthy control) X condition (stress, control) mixed model ANOVA, corrected for age, gender and the order of sessions (i.e. stress induction at first or second session). Post-hoc analyses were performed in case of significance to determine the direction of the interaction effect. In addition, we analyzed group differences during the control visit with a two sample t-test to assess whether patients showed altered connectivity without the presence of stress, again corrected for age, gender and the order of sessions. Voxel-wise statistical tests were family wise error (FWE) rate corrected (two-sided; P<0.05) for multiple comparisons across the whole brain at cluster level using an initial height threshold of P<0.001 with a small volume correction for the regions targeted by our seeds. Because we were most interested in connectivity within both hypothesized networks, we performed a small volume correction for the regions targeted by our seeds, i.e. the supplementary motor area (SMA) for the posterior putamen and the ventromedial prefrontal cortex (vmPFC) for the caudate nucleus. We used the WFU Pickatlas toolbox in SPM12 to define the SMA ROI according to the anatomical automatic labeling (AAL) atlas. For the goal-directed ROI, we selected coordinates of the vmPFC that have shown to be active during goal-directed behavior in an instrumental discrimination task using a sphere with 10mm radius [33]. Movement during scanning was investigated by calculating the mean framewise displacement (FD) for each scanning session, and was compared between groups with a Mann-Whitney U test. To further investigate whether the imaging results were linked to the severity and type of symptoms, ratings on the obsession and compulsion scale of the YBOCS were correlated with the connectivity results extracted from the significant clusters using a Pearson correlation in SPSS (see Supplementary Information).

## Results

### Demographic and clinical data

Demographic and clinical data are presented in Table 1. There were no significant group differences in age, sex distribution, level of education and movement during scanning. As expected, ratings on the Y-BOCS, HAM-A and HDRS were significantly higher in patients than in controls. Patients had various current comorbid disorders, including panic disorder (n=2), agoraphobia (n=1), hypochondria (n=1), generalized anxiety disorder (n=1), social anxiety disorder (n=1), past major depressive disorder (n=6) and past alcohol abuse (n=1). In addition, three patients matched the criteria for a personality disorder, i.e. obsessive-compulsive, antisocial and avoidant personality disorder. Furthermore, nineteen patients used a stabile dose of medication during the two visits, including fluoxetine, (es)citalopram, sertraline, paroxetine, clomipramine or venlafaxine.

**TABLE 1;.** Demographic and clinical data presented as the mean ± SD.

|  | Patients (n=23) | Controls (n=23) | P-value |
| --- | --- | --- | --- |
| Age (years) | 33.48 ± 2.02 | 33.52 ± 3.07 | p=0.605<sup>1</sup> |
| Gender (male/female) | 10/13 | 12/11 | p=0.555<sup>2</sup> |
| Handedness (left/right) | 2/21 | 4/19 | p=0.381<sup>2</sup> |
| Highest education (n) | 3/5/7/3/5 | 0/4/10/4/5 | p=0.436<sup>2</sup> |
| – lower professional | 3 | 0 |  |
| – secondary school | 5 | 4 |  |
| – medium professional | 7 | 10 |  |
| – higher professional | 8 | 9 |  |
| Y-BOCS: |  |  |  |
| – total | 22.57 ± 1.24 | 0.13 ± 0.10 | p<0.001<sup>1*</sup> |
| – obsessions | 11.61 ± 0.64 | 0.13 ± 0.10 | |
| – compulsions | 10.52 ± 0.84 | 0.00 ± 0.00 | |
| HAM-A | 11.83 ± 1.57 | 1.39 ± 1.64 | p<0.001<sup>1*</sup> |
| HDRS | 8.17 ± 1.24 | 0.65 ± 0.19 | p<0.001<sup>1*</sup> |
| Medication status (yes/no) | 19/4 | 0/23 |  |
| Framewise displacement (mm) |  |  |  |
| – control visit | 0.14 ± 0.02 | 0.12 ± 0.01 | p=0.538<sup>1</sup> |
| – stress visit | 0.16 ± 0.03 | 0.12 ± 0.01 | p=0.070<sup>1</sup> |

\* Significant difference between groups, $p \leq 0.05$ , tested with <sup>1</sup> Mann-Whitney U test or <sup>2</sup> chi-square test. Scales: Yale-Brown Obsessive-Compulsive Scale (Y-BOCS), Hamilton Anxiety Rating Scale (HAM-A), Hamilton Depression Rating Scale (HDRS).

### Stress response

Physiological and subjective stress responses are presented in Figure 2. Blood pressure and heart rate were analyzed using a mixed model ANOVA with the factors group (patients, healthy controls), condition (stress, control) and time (five time points), corrected for the order of sessions. For systolic blood pressure, this analysis revealed a main effect of condition with higher systolic blood pressure in the stress condition (F(1,35)=4.64, p=0.038, η_p_^2^=0.12), a main effect of time (F(4,140) = 3.20, p=0.015, η_p_^2^=0.08) and a condition X time interaction (F(4,140)=8.91, p<0.001, η_p_^2^=0.20). The condition X time interaction was due to higher systolic blood pressure after the math task during the stress condition compared to the control condition across groups (t(45)=6.35, p<0.001). The analysis of the diastolic blood pressure revealed a main effect of condition with higher diastolic blood pressure in the stress condition (F(1,34)=4.24, p=0.047, η_p_^2^=0.11). As expected, the analysis of the heart rate did not show any significant results [27]. For cortisol, the analysis of baseline level showed no significant effects. We then calculated the AUCg and entered these values into an interaction analysis with the factors group (patients, healthy controls) and condition (stress, control). The analysis revealed a group X condition interaction (F(1,43)=6.36, p=0.015, η_p_^2^=0.13). Post-hoc testing showed significantly higher cortisol levels in the stress condition than the control condition in healthy controls (t(22)=3.83, p=0.001) but not in patients. The analysis of subjective anxiety levels, as measured with the STAI, revealed a main effect of condition with higher anxiety levels in the stress condition (F(1,40)=5.20, p=0.028, η_p_^2^=0.12), a main effect of time (F(2.08,83.12)=4.45, p=0.014, η_p_^2^=0.10) and a condition X time interaction (F(2.61,104.28)=12.31, p<0.001, η_p_^2^=0.24). In addition, there was a significant main effect of group (F(1,40)=46.3, p<0.001, η_p_^2^=0.54), indicating that patients reported higher anxiety levels than healthy controls overall. Post-hoc paired T-testing showed that all participants reported significantly higher anxiety levels in the stress condition compared to the control condition after the stress procedure (t(45) = 7.17, p<0.001), with a trend towards significance during scanning (t(43) = 2.22, p=0.032). The analysis of the subjective stress level questionnaire showed a main effect of condition with higher ratings in the stress condition (F(1,42)=105.99, p<0.001, η_p_^2^=0.72). Analyzing the specific subscales showed that all participants rated the stress condition as more difficult (t(44)=12.09, p<0.001), more unpleasant (t(44)=13.02, p<0.001), and more stressful (t(44)=10.8, p<0.001) than the control condition. Together, these results show that the stress condition successfully increased blood pressure and psychological stress measures in both groups, and that group differences were only present for the stress-induced cortisol response, and anxiety levels (independent of stress induction).

**FIGURE 2;.**
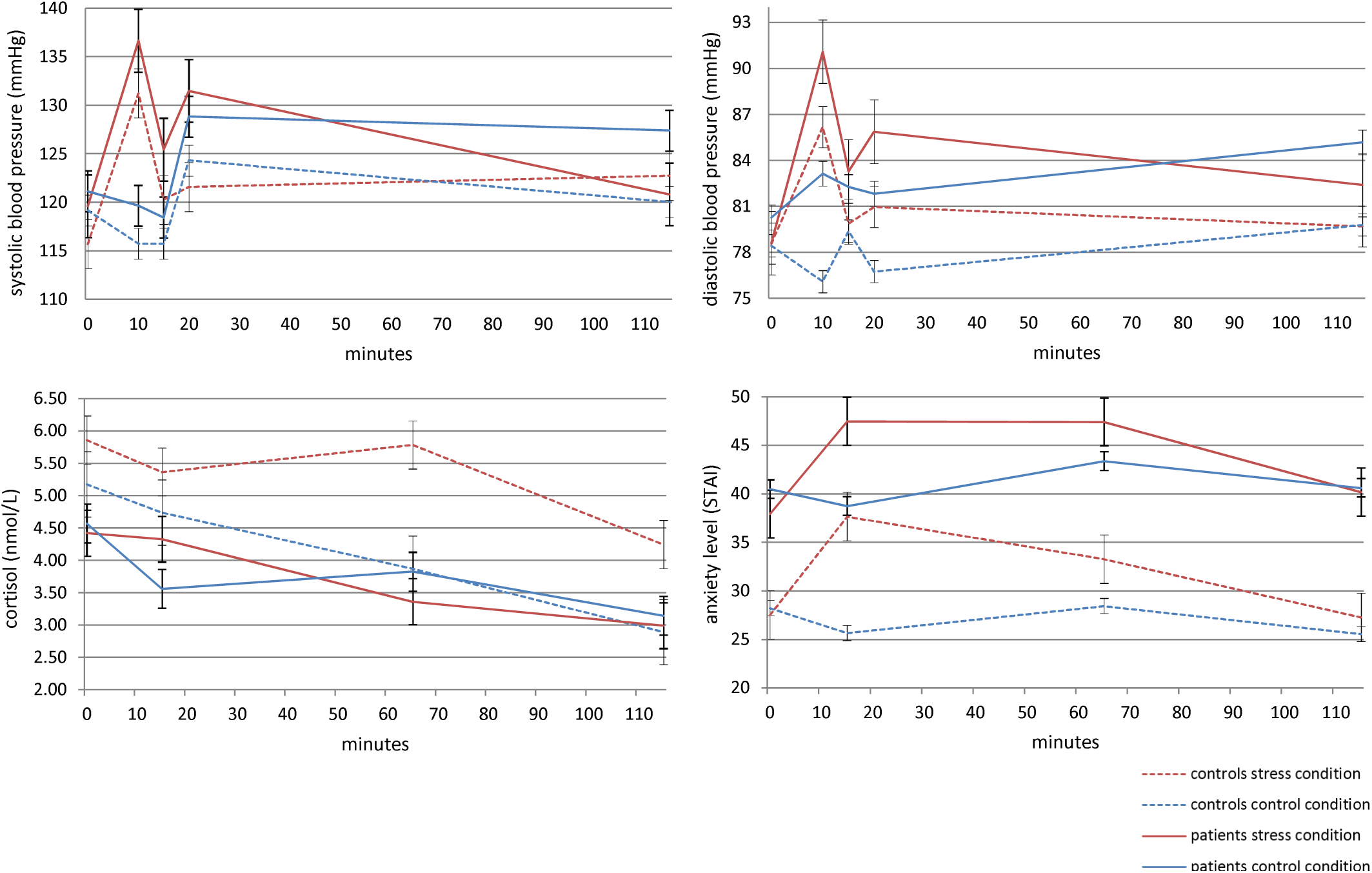
Physiological and subjective stress response, time points: baseline (0 min), after water test (10 min), after math task (15 min), before scanning session (20 min), before resting-state scan (65 min) and after scanning (115 min) reported as the mean with standard error.

### Seed-to-voxel analysis

To assess the effects of stress induction on these networks, we performed a mixed model ANOVA with the factors group (patients, healthy controls) and condition (stress, control), corrected for the order of sessions. This analysis revealed a significant interaction for the goal-directed seeds (for significant clusters, see Table 2). Compared to healthy controls, patients showed a blunted decrease in connectivity between the left and right caudate nuclei and precuneus after stress induction (Figure 3). This interaction effect was mainly driven by decreased connectivity after stress in healthy controls between the right caudate nucleus and precuneus and the left caudate nucleus and precuneus. The increase in connectivity after stress in patients was not significant. We found no significant clusters in the vmPFC using a small volume correction. For the habit seeds, we found no significant group differences in connectivity after stress between the posterior putamen and the SMA, nor with other regions of the brain. Furthermore, there was no main effect of stress across patients and controls in any of the seeds.

**TABLE 2;.** Significant clusters from the seed-to-voxel group X condition interaction analysis.

| Seed | MNI coordinates |  |  | Cluster size | P-value | Region | BA |
| --- | --- | --- | --- | --- | --- | --- | --- |
|  | x | y | z |  |  |  |  |
| OCD patients > healthy controls (stress > control condition) |  |  |  |  |  |  |  |
| Caudate nucleus L | -6 | -62 | +38 | 92 | 0.038 | precuneus | 7 |
| Caudate nucleus R | -4 | -66 | +42 | 467 | <0.001 | precuneus | 7 |
| Posterior putamen L |  |  |  |  | no clusters |  |  |
| Posterior putamen R |  |  |  |  | no clusters |  |  |

**FIGURE 3;.**
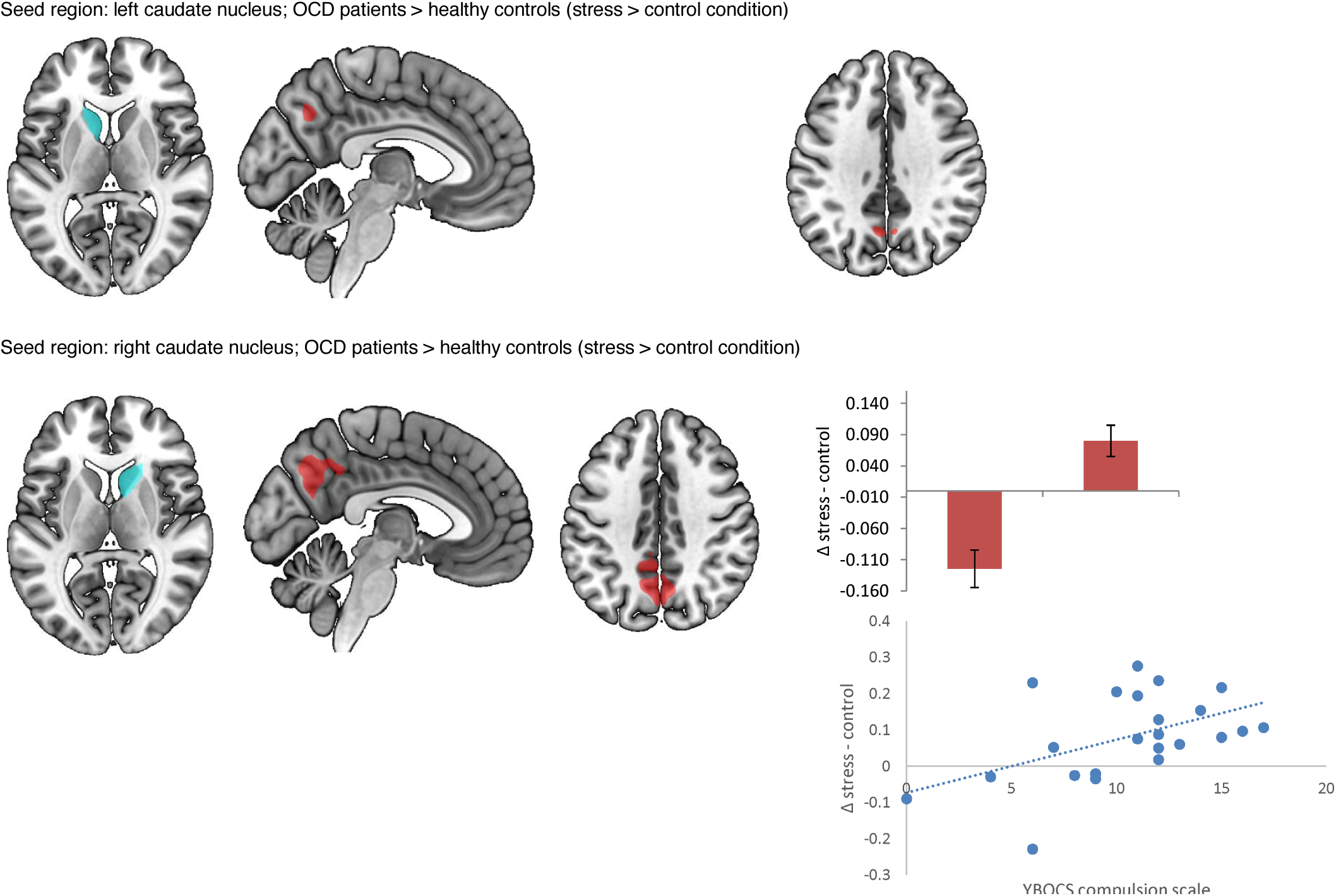
Stress effects in patients compared to healthy controls with the selected seeds, significant connectivity results (thresholded at p>0.001 for illustrational purposes), contrast estimates extracted from the significant clusters, and correlation with compulsion scale of the YBOCS.

We further investigated whether the connectivity results were linked to the severity and type of symptoms, by correlating the extracted values from the significant clusters of the stress versus control contrast with the ratings on the obsession and compulsion scale of the Y-BOCS in patients only. This showed a significant correlation between the connectivity of the right caudate nucleus and precuneus and ratings on the compulsion scale of the Y-BOCS (r=0.49, n=23, p=0.017, see Figure 3), indicating that patients who are more compulsive show a larger increase in connectivity in the aftermath of stress. There were no significant correlations between the imaging results and the rating on the obsession scale of the Y-BOCS. We then analyzed group differences during the control session with a two sample t-test to see whether patients showed altered connectivity without the presence of stress, corrected for age, gender and the order of sessions. Compared to healthy controls, patients showed significantly lower connectivity between the caudate nucleus and lateral occipital cortex, the angular gyrus, the middle and superior frontal gyrus, the precuneus cortex, the posterior cingulate gyrus and the frontal pole (p(FWE)<0.05, for significant clusters see Supplemental Information Table 1. Both the left and right posterior putamen showed no significant differences in baseline connectivity between patients and controls.

## Discussion

This is the first study investigating the effects of stress on intrinsic functional connectivity in patients with OCD, which aimed to test the hypothesis that stress induces a shift from the goal-directed towards the habit network. Using a seed-to-voxel resting-state analysis, we found a blunted reduction in connectivity between the seeds involved in the goal-directed network (i.e. the left and right caudate nucleus) and the precuneus in patients with OCD. This result was positively correlated with the severity of compulsive symptoms. Stress did not have an effect on functional connectivity between the seeds involved in the habit network (i.e. the left and right posterior putamen) and the rest of the brain, including the supplementary motor areas. Measures of blood pressure and subjective anxiety confirmed successful stress induction across groups, and showed significantly lower cortisol responses in patients compared to healthy controls. Together, these results suggest that stress-induced compulsivity is primarily associated with recruitment of the goal-directed network, rather than with changes in the habit network. The habit theory of OCD is centered around the balance between the goal-directed and habit networks, but ignores the influence of distress. Distress however, is a central aspect of the cognitive-behavioral theory of OCD. Although we found no evidence that stress leads to an overdependence on the habit network, the result of altered goal-directed connectivity does link the apparent contradictory theories of OCD and suggest that both theories contribute to the pathophysiology of OCD.

Our results were not fully in line with our hypothesis, as stress did not affect intrinsic connectivity within the habit network, and we did not observe reduced connectivity of the goal-directed network with the ventromedial prefrontal cortex. Instead, the results point towards altered connectivity between the caudate nucleus and the precuneus, an area that is part of the default mode network (DMN). The precuneus is involved in self-consciousness and self-referential processing, defined as the process of relating information to the self [34,35]. When individuals are not involved in an attention demanding or goal-directed task, the DMN is activated and self-referential processing is believed to predominate [36]. Compared to healthy controls, patients showed decreased baseline connectivity between the goal-directed network and the precuneus. After stress, healthy controls showed uncoupling of the goal-directed network and DMN, which implies that stress normally induces upscaling of the goal-directed network at the cost of the DMN. In contrast, OCD patients did not show a decrease in connectivity between the caudate nucleus and precuneus after stress, suggesting that patients might be stuck in self-referential processing at the cost of goal-directed behavior. Almost all OCD symptoms can be narrowed down to the underlying desire for absolute certainty and maintaining control, therefore obsessions usually develop in domains where there is no absolute level of certainty, such as sexuality, sickness and danger [3]. Unfortunately patients do not base the level of certainty on external reality, but on feelings of anxiousness and unrealistic thoughts. This causes them to continuously focus on themselves, resulting in a mental loop of self-referential processing. The lack of downscaling of the DMN that was seen in patients might thus be the result of compulsive self-referential processing under stress. The stress-induced connectivity between the right caudate nucleus and the precuneus was positively correlated with the compulsion scale of the Y-BOCS, indicating that patients who are prone to compulsive behavior are more likely to show a blunted reduction in connectivity after stress. This further confirms the assumption that our results can be interpreted as stress-induced self-referential processing at the cost of goal-directed control, resulting in mental compulsions.

Remarkably, we found no stress induced coupling between the posterior putamen and SMA. This leads to the speculation that compulsions are more strongly related to impaired goal-directed control than to overdependence on habits. Our null finding could be due to the study design, as the motor circuit is not probed during resting-state. Future research is needed to see whether these results are confirmed using a task that requires goal-directed and habitual behavior for further understanding of the pathophysiology of OCD. Furthermore, we cannot rule out that there were early effects of stress that are missed because of the delay between the stress induction and the resting-state scan. This study was part of a larger trial, therefore the resting-state scan was conducted approximately 65 minutes after the SECPT. Although patients were no longer in the acute stress state, cortisol levels in the control group were still elevated (Figure 2), and research has shown that the effects of corticosteroids can continue for several hours [19]. Additionally, participants still rated their anxiety level to be higher before the resting-state scan started in the stress visit compared to the control visit, implicating that the stress response was prolonged during the scanning session.

In addition to the group differences in connectivity, patients also showed lower stress-induced cortisol levels compared to healthy controls. This blunted cortisol response could be due to adaptation of stress circuits as a consequence of chronic exposure to stress. This mechanism is seen across multiple psychiatric disorders including post-traumatic stress disorder, major depressive disorder and panic disorder [37]. Similarly, previous studies have also shown a non-response of cortisol during stressful exposure and response prevention therapy in OCD [38,39].

Without the presence of stress, the goal-directed seeds showed extensive group differences in connectivity with frontal areas, whereas the habit seeds showed no differences in connectivity. This is consistent with the hypothesis that patients already show an impairment in the goal-directed network, regardless of the presence of induced stress [4]. The finding of reduced connectivity between the caudate and these cortical regions is inconsistent with corticostriatal hyperconnectivity that is often described in OCD [40-42], but has previously been demonstrated in research conducted in unmedicated patients [43]. This study has several limitations. Resting-state fMRI is a promising technique for investigating the functional architecture of the brain, but the results provide no information on the direction of the connectivity. Therefore, we can only speculate on causal relationships between significant nodes [44]. As mentioned above, we might have missed stress-induced effects within the habit network in the acute stress phase due to the delay between the stress induction and the resting-state scan. Last, the majority of our patients were using medication and had various comorbid disorders which could have influenced the imaging results. Despite these limitations, our results show that stress influences functional connectivity of the goal-directed network in OCD. These findings provide first insight into the late effects of stress on the balance between the goal-directed and habit network in OCD, implicating that stress-induced compulsivity is predominantly associated with involvement of the goal-directed network.

## Acknowledgements

This work is supported by ZonMw grants no. 165.610.002 and 016.156.318.

## References

1. Ruscio AM, Stein DJ, Chiu WT, Kessler RC. The epidemiology of obsessive-compulsive disorder in the National Comorbidity Survey Replication. Mol Psychiatry. 2010;15(1):53–63.

2. Salkovskis PM. Obsessional-compulsive problems: a cognitive-behavioural analysis. Behav Res Ther. 1985;23(5):571–83.

3. Denys D. Obsessionality & compulsivity: a phenomenology of obsessive-compulsive disorder. Philos Ethics Humanit Med. 2011;6:3.

4. Gillan CM, Papmeyer M, Morein-Zamir S, Sahakian BJ, Fineberg NA, Robbins TW, et al. Disruption in the balance between goal-directed behavior and habit learning in obsessivecompulsive disorder. Am J Psychiatry. 2011;168(7):718–26.

5. Gillan CM, Morein-Zamir S, Urcelay GP, Sule A, Voon V, Apergis-Schoute AM, et al. Enhanced avoidance habits in obsessive-compulsive disorder. Biol Psychiatry. 2014;75(8):631–8.

6. Gillan CM, Robbins TW, Sahakian BJ, van den Heuvel OA, van Wingen G. The role of habit in compulsivity. Eur Neuropsychopharmacol. 2016;26(5):828–40.

7. Balleine BW, O’Doherty JP. Human and rodent homologies in action control: corticostriatal determinants of goal-directed and habitual action. Neuropsychopharmacology. 2010;35(1):48–69.

8. de Wit S, Watson P, Harsay HA, Cohen MX, van de Vijver I, Ridderinkhof KR. Corticostriatal connectivity underlies individual differences in the balance between habitual and goal-directed action control. J Neurosci. 2012;32(35):12066–75.

9. Alexander GE, Crutcher MD, DeLong MR. Basal ganglia-thalamocortical circuits: parallel substrates for motor, oculomotor, “prefrontal” and “limbic” functions. Prog Brain Res. 1990;85:119–46.

10. Watson P, van Wingen G, de Wit S. Conflicted between Goal-Directed and Habitual Control, an fMRI Investigation. eNeuro. 2018;5(4).

11. Radua J, van den Heuvel OA, Surguladze S, Mataix-Cols D. Meta-analytical comparison of voxelbased morphometry studies in obsessive-compulsive disorder vs other anxiety disorders. Arch Gen Psychiatry. 2010;67(7):701–11.

12. Milad MR, Rauch SL. Obsessive-compulsive disorder: beyond segregated cortico-striatal pathways. Trends Cogn Sci. 2012;16(1):43–51.

13. American Psychiatric Association. Diagnostic and statistical manual of mental disorders (5th ed.). American Psychiatric Publishing: Arlington, VA; 2013.

14. Rachman S. A cognitive theory of obsessions: elaborations. Behav Res Ther. 1998;36(4):385–401.

15. Schwabe L, Wolf OT. Stress prompts habit behavior in humans. J Neurosci. 2009;29(22):7191–8.

16. Otto AR, Raio CM, Chiang A, Phelps EA, Daw ND. Working-memory capacity protects modelbased learning from stress. Proc Natl Acad Sci U S A. 2013;110(52):20941–6.

17. Adams TG, Kelmendi B, Brake CA, Gruner P, Badour CL, Pittenger C. The role of stress in the pathogenesis and maintenance of obsessive-compulsive disorder. Chronic Stress (Thousand Oaks). 2018;2.

18. de Kloet ER, Joels M, Holsboer F. Stress and the brain: from adaptation to disease. Nat Rev Neurosci. 2005;6(6):463–75.

19. Hermans EJ, Henckens MJ, Joels M, Fernandez G. Dynamic adaptation of large-scale brain networks in response to acute stressors. Trends Neurosci. 2014;37(6):304–14.

20. Henckens MJ, van Wingen GA, Joels M, Fernandez G. Time-dependent corticosteroid modulation of prefrontal working memory processing. Proc Natl Acad Sci U S A. 2011;108(14):5801–6.

21. Henckens MJ, van Wingen GA, Joels M, Fernandez G. Time-dependent effects of corticosteroids on human amygdala processing. J Neurosci. 2010;30(38):12725–32.

22. Hamilton M. The assessment of anxiety states by rating. Br J Med Psychol. 1959;32(1):50–5.

23. Hamilton M. A rating scale for depression. J Neurol Neurosurg Psychiatry. 1960;23:56–62.

24. Goodman WK, Price LH, Rasmussen SA, Mazure C, Fleischmann RL, Hill CL, et al. The Yale-Brown Obsessive Compulsive Scale. I. Development, use, and reliability. Arch Gen Psychiatry. 1989;46(11):1006–11.

25. Sheehan DV, Lecrubier Y, Sheehan KH, Amorim P, Janavs J, Weiller E, et al. The Mini-International Neuropsychiatric Interview (M.I.N.I.): the development and validation of a structured diagnostic psychiatric interview for DSM-IV and ICD-10. J Clin Psychiatry. 1998;59 Suppl 20:22–33;quiz 34-57.

26. Pfohl BB, N.; Zimmerman, M.; Stangl, D.;. Structured interview for DSM-IV personality (SIDP-IV). American Psychiatric Association: Washington (DC); 1997.

27. Schwabe L, Haddad L, Schachinger H. HPA axis activation by a socially evaluated cold-pressor test. Psychoneuroendocrinology. 2008;33(6):890–5.

28. D Spielberger C, Gorsuch R, E Lushene R, Vagg PR, A Jacobs G. Manual for the State-Trait Anxiety Inventory (Form Y1 – Y2). 1983.

29. Poser BA, Versluis MJ, Hoogduin JM, Norris DG. BOLD contrast sensitivity enhancement and artifact reduction with multiecho EPI: parallel-acquired inhomogeneity-desensitized fMRI. Magn Reson Med. 2006;55(6):1227–35.

30. Pruessner JC, Kirschbaum C, Meinlschmid G, Hellhammer DH. Two formulas for computation of the area under the curve represent measures of total hormone concentration versus time-dependent change. Psychoneuroendocrinology. 2003;28(7):916–31.

31. Behzadi Y, Restom K, Liau J, Liu TT. A component based noise correction method (CompCor) for BOLD and perfusion based fMRI. Neuroimage. 2007;37(1):90–101.

32. Whitfield-Gabrieli S, Nieto-Castanon A. Conn: a functional connectivity toolbox for correlated and anticorrelated brain networks. Brain Connect. 2012;2(3):125–41.

33. de Wit S, Corlett PR, Aitken MR, Dickinson A, Fletcher PC. Differential engagement of the ventromedial prefrontal cortex by goal-directed and habitual behavior toward food pictures in humans. J Neurosci. 2009;29(36):11330–8.

34. Cavanna AE, Trimble MR. The precuneus: a review of its functional anatomy and behavioural correlates. Brain. 2006;129(Pt 3):564–83.

35. Northoff G, Heinzel A, Greck M, Bennpohl F, Dobrowolny H, Panksepp J. Self-referential processing in our brain - A meta-analysis of imaging studies on the self. Neuroimage. 2006;31(1):440–57.

36. Buckner RL, Andrews-Hanna JR, Schacter DL. The brain’s default network: anatomy, function, and relevance to disease. Ann N Y Acad Sci. 2008;1124:1–38.

37. Wichmann S, Kirschbaum C, Bohme C, Petrowski K. Cortisol stress response in post-traumatic stress disorder, panic disorder, and major depressive disorder patients. Psychoneuroendocrinology. 2017;83:135–41.

38. Gustafsson PE, Gustafsson PA, Ivarsson T, Nelson N. Diurnal cortisol levels and cortisol response in youths with obsessive-compulsive disorder. Neuropsychobiology. 2008;57(1-2):14–21.

39. Kellner M, Wiedemann K, Yassouridis A, Muhtz C. Non-response of cortisol during stressful exposure therapy in patients with obsessive-compulsive disorder--preliminary results. Psychiatry Res. 2012;199(2):111–4.

40. Harrison BJ, Soriano-Mas C, Pujol J, Ortiz H, Lopez-Sola M, Hernandez-Ribas R, et al. Altered corticostriatal functional connectivity in obsessive-compulsive disorder. Arch Gen Psychiatry. 2009;66(11):1189–200.

41. Harrison BJ, Pujol J, Cardoner N, Deus J, Alonso P, Lopez-Sola M, et al. Brain corticostriatal systems and the major clinical symptom dimensions of obsessive-compulsive disorder. Biol Psychiatry. 2013;73(4):321–8.

42. Fitzgerald KD, Welsh RC, Stern ER, Angstadt M, Hanna GL, Abelson JL, et al. Developmental alterations of frontal-striatal-thalamic connectivity in obsessive-compulsive disorder. J Am Acad Child Adolesc Psychiatry. 2011;50(9):938–48 e3.

43. Posner J, Marsh R, Maia TV, Peterson BS, Gruber A, Simpson HB. Reduced functional connectivity within the limbic cortico-striato-thalamo-cortical loop in unmedicated adults with obsessive-compulsive disorder. Hum Brain Mapp. 2014;35(6):2852–60.

44. Cole DM, Smith SM, Beckmann CF. Advances and pitfalls in the analysis and interpretation of resting-state FMRI data. Front Syst Neurosci. 2010;4:8.

