## Supplemental Information for "The effect of stress on the balance between goal-directed and habit networks in obsessive-compulsive disorder"

**Supplemental methods**

*Exclusion criteria*

Exclusion criteria were (1) having a current major depressive disorder, bipolar disorder, psychotic disorder or alcohol or substance dependence as assessed with the Mini-International Neuropsychiatric Interview (M.I.N.I. [1]) (2) having a history of major head trauma or neurological disease; (3) any MRI contraindications; (4) a self-reported inability or unease to stop smoking 3 hours prior to testing; (5) having any endocrinological disorder or regularly using corticosteroids; (6) undergoing a current treatment with antipsychotic medication; (7) using benzodiazepines or recreational drugs 72 hours prior to each test session or using alcohol within 24 hours before each measurement; and (8) having an irregular sleep/wake rhythm.

*Socially evaluated cold-pressor test (SECPT)*

First, participants had to hold their right hand and wrist in ice water (1-3°C) for three minutes while being videotaped and monitored by an unfamiliar investigator. This unfamiliar person was instructed to wear a white lab coat and act unsympathetically, thereby inducing social stress. Afterwards, participants were asked to count backwards from 2059 in steps of 17. In case the participant made a mistake they were told to start over. During the control session, participants had to immerse their right hand in warm water (35-37°C) without being videotaped or monitored by an unfamiliar investigator. Following the warm water procedure, they were asked to count to 600 in steps of 10.

*Data acquisition*

MR imaging was performed on a 3.0 T Philips MRI scanner (Philips, Best, The Netherlands), using a 32-channel SENSE head coil. The scanning protocol included a high-resolution T1-weighted MRI (sequence parameters: repetition time=6.9ms; echo time=3.1ms; flip angle=8°; 150 sagittal slices; voxel size=1.1mm isotropic, total duration=179s). For fMRI, multi-echo echoplanar imaging (EPI) was used to acquire T2*-weighted MRI volumes with blood oxygen level-dependent (BOLD) contrast (sequence parameters: repetition time=2375ms; number of echos=3, echo time=9, 26 and 44ms; flip angle=76°; field of view=224mm x 122mm x 224mm; voxel size=3.0 mm isotropic; 37 slices; 200 volumes with a total duration of 8 minutes). Multi-echo EPI reduces signal dropout and distortion while enhancing BOLD contrast sensitivity [2]. Three dummy scans were acquired to allow for equilibration of the signal. Head immobilization was established using foam pads inside the coil. Participants were instructed to stay awake and relax with their eyes open while thinking of anything that came to their mind.

*MRI analysis*

Functional MRI data were preprocessed using SPM12 (www.fil.ion.ucl.ac.uk/spm/) and the CONN toolbox v18 (http://www.nitrc.org/projects/conn) in Matlab version 2014b (http://www.mathworks.com). First, realignment was performed and the three echoes were combined using in-house software to optimize BOLD contrast sensitivity and minimize signal dropout and distortion [2]. This was followed by slice-time correction. Artifacts were detected for scrubbing using the ART-based functional outlier detection. Volumes were considered outliers if they exceeded the global z signal threshold of 9 or the motion threshold of 2mm. Following co-registration of the mean EPI to the T1-weighted structural scan, functional images were normalized to Montreal Neurological Institute (MNI) space and smoothed with a 3D Gaussian kernel of 6mm at FWHM. Participants with realignment parameters exceeding 3mm translation on the x-, y-, or z-axis were excluded from the analysis, resulting in the exclusion of two patients. One healthy control was excluded due to an incidental neurological finding and one control was excluded because of insufficient brain coverage due to inaccurate setting of the field of view. This led to the inclusion of twenty-three patients and twenty-three healthy controls for further analysis.

After preprocessing, six principal components were derived from white matter and ventricles and they were included as nuisance parameters using CompCor [3]. Motion-related artefacts were accounted for by regressing out realignment parameters. In addition, volumes that exceeded the threshold of global signal change (i.e. Z-value=9) or motion (i.e. 2mm) were considered an outlier and were censored using scrubbing. Based on our hypothesis, we selected four seeds (i.e. left and right caudate for the goal-directed network, left and right posterior putamen for the habit network) for further analysis using the Harvard-Oxford atlas. The putamen was delineated at y=2 as described in previous research [4], resulting in the region y≤2 for the posterior putamen. Because the caudate nucleus and posterior putamen are located closely to each other, the timeseries from the ipsilateral seed were regressed out to ensure that the results were specific for the selected seeds and did not reflect striatal connectivity in general. The resulting residual time series were then band-pass filtered at 0.008-0.09 Hz and were used to create individual connectivity maps through bivariate correlation and Fisher’s r-to-z transformation [5]. We performed a seed-to-voxel analysis for the selected seeds (i.e. left and right caudate, left and right posterior putamen) to investigate whether there were group differences in stress-induced connectivity. Connectivity maps were entered into a second-level group (patient, healthy control) X condition (stress, control) mixed model ANOVA, corrected for age, gender and the order of sessions (i.e. stress induction at first or second session). Post-hoc analyses were performed in case of significance to determine the direction of the interaction effect. In addition, we analyzed group differences during the control visit with a two sample T-test to assess whether patients showed altered connectivity without the presence of stress, again corrected for age, gender and the order of sessions. Voxel-wise statistical tests were family wise error (FWE) rate corrected (two-sided; P<0.05) for multiple comparisons across the whole brain at cluster level using an initial height threshold of P<0.001. Because we were most interested in connectivity within both hypothesized networks, we performed a small volume correction for the regions targeted by our seeds, i.e. the supplementary motor area (SMA) for the posterior putamen and the ventromedial prefrontal cortex (vmPFC) for the caudate nucleus. We used the WFU Pickatlas toolbox in SPM12 to define the SMA ROI according to the anatomical automatic labeling (AAL) atlas. For the goal-directed ROI, we selected coordinates of the vmPFC that have shown to be active during goal-directed behavior in an instrumental discrimination task using a sphere with 10mm radius [4]. Movement during scanning was investigated by calculating the mean framewise displacement (FD) for each scanning session, and was compared between groups with a Mann-Whitney U test. To further investigate whether the imaging results were linked to the severity and type of symptoms, ratings on the obsession and compulsion scale of the YBOCS were correlated with the connectivity results extracted from the significant clusters using a Pearson correlation in SPSS.

**References**

1	Sheehan DV, Lecrubier Y, Sheehan KH, Amorim P, Janavs J, Weiller E, et al. The Mini-International Neuropsychiatric Interview (M.I.N.I.): the development and validation of a structured diagnostic psychiatric interview for DSM-IV and ICD-10. J Clin Psychiatry. 1998;59 Suppl 20:22-33;quiz 34-57.

2	Poser BA, Versluis MJ, Hoogduin JM, Norris DG. BOLD contrast sensitivity enhancement and artifact reduction with multiecho EPI: parallel-acquired inhomogeneity-desensitized fMRI. Magn Reson Med. 2006;55(6):1227-35.

3	Behzadi Y, Restom K, Liau J, Liu TT. A component based noise correction method (CompCor) for BOLD and perfusion based fMRI. Neuroimage. 2007;37(1):90-101.

4	de Wit S, Corlett PR, Aitken MR, Dickinson A, Fletcher PC. Differential engagement of the ventromedial prefrontal cortex by goal-directed and habitual behavior toward food pictures in humans. J Neurosci. 2009;29(36):11330-8.

5	Whitfield-Gabrieli S, Nieto-Castanon A. Conn: a functional connectivity toolbox for correlated and anticorrelated brain networks. Brain Connect. 2012;2(3):125-41.
